# Ceftazidime Is a Potential Drug to Inhibit SARS-CoV-2 Infection In Vitro by Blocking Spike Protein-ACE2 Interaction

**DOI:** 10.1101/2020.09.14.295956

**Authors:** ChangDong Lin, Yue Li, MengYa Yuan, MengWen Huang, Cui Liu, Hui Du, XingChao Pan, YaTing Wen, Xinyi Xu, Chenqi Xu, JianFeng Chen

## Abstract

Coronavirus Disease 2019 (COVID-19) spreads globally as a sever pandemic, which is caused by severe acute respiratory syndrome coronavirus 2 (SARS-CoV-2). Cell entry of SARS-CoV-2 mainly depends on binding of the viral spike (S) proteins to angiotensin converting enzyme 2 (ACE2) on host cells. Therefore, repurposing of known drugs to inhibit S protein-ACE2 interaction could be a quick way to develop effective therapy for COVID-19. Using a high-throughput screening system to investigate the interaction between spike receptor binding domain (S-RBD) and ACE2 extracellular domain, we screened 3581 FDA-approved drugs and natural small molecules and identified ceftazidime as a potent compound to inhibit S-RBD–ACE2 interaction by binding to S-RBD. In addition to significantly inhibit S-RBD binding to HPAEpiC cells, ceftazidime efficiently prevented SARS-CoV-2 pseudovirus to infect ACE2-expressing 293T cells. The inhibitory concentration (IC_50_) was 113.2 μM, which is far below the blood concentration (over 300 μM) of ceftazidime in patients when clinically treated with recommended dose. Notably, ceftazidime is a drug clinically used for the treatment of pneumonia with minimal side effects compared with other antiviral drugs. Thus, ceftazidime has both anti-bacterial and anti-SARS-CoV-2 effects, which should be the first-line antibiotics used for the clinical treatment of COVID-19.

## INTRODUCTION

Coronavirus disease 2019 (COVID-19) has spread globally as a severe pandemic, which was caused by a novel coronavirus named Severe Acute Respiratory Syndrome coronavirus 2 (SARS-CoV-2) ^1^. According to the latest statistics of COVID-19 released by Johns Hopkins University on September 1, 2020, there were 25.50 million confirmed cases and 0.85 million deaths globally. COVID-19 has become a serious threat to human survival and is likely to coexist with human beings for a long time. Unfortunately, there is still no effective cure for COVID-19, especially the critically ill patients.

Similar to SARS-CoV, SARS-CoV-2 belongs to the sarbecoviruses (Betacoronirus, Coronaviridae) and can cause life-threatening respiratory diseases ^2–4^. Some mechanisms underlying SARS-CoV-2 infection of target cells have been reported ^5–7^. Spike (S) protein on the surface of SARS-CoV-2 facilitates viral entry into target cells by mediating virus receptor recognition and membrane fusion. The N-terminal region of its S1 domain contains receptor binding domain (RBD), which directly binds to angiotensin converting enzyme 2 (ACE2) receptor on the plasma membrane of host cells and is responsible for virus attachment ^8,9^. Therefore, blocking the binding of spike protein to ACE2 is an effective way to inhibit the infection of target cells by SARS-CoV-2. By now, several studies have reported the development of monoclonal antibodies targeting spike protein ^10,11^, however, the typical timeline for approval of a novel antibody for the management viral infection is long. In addition, the side effects such as antibody-dependent enhancement of viral infection need to be considered ^11–13^, and the high cost of antibody treatment will limit the clinical application. Therefore, repurposing of known small molecule drugs to inhibit spike protein and ACE2 binding could significantly accelerate the deployment of effective and affordable therapies for COVID-19.

In this study, we expressed and purified Spike-RBD (S-RBD) and the extracellular domain of ACE2 (ACE2-ECD) and then established an AlphaScreen-based high-throughput system for screening small molecules that block S-RBD–ACE2-ECD interaction. From 3581 Food and Drug Administration (FDA)-approved drugs and natural small molecule compounds, we identified 10 compounds that could block S-RBD–ACE2-ECD interaction in the initial screening. Notably, ceftazidime bound to S-RBD and showed the strongest potency for the inhibition of S-RBD binding to human pulmonary alveolar epithelial cells (HPAEpiC). Moreover, ceftazidime efficiently inhibited the infection of ACE2-expressing 293T cells by SARS-CoV-2 pseudovirus. Overall, we identified ceftazidime as a potential drug to inhibit SARS-CoV-2 infection with minimal known side effects and affordable price.

## RESULTS

### Establishment of AlphaScreen system to detect S-RBD–ACE2 interaction

In order to screen small molecules that block S protein-ACE2 binding, we firstly established an AlphaScreen-based high-throughput system to detect the interaction between S-RBD and ACE2-ECD (Fig. 1a). S-RBD and ACE2-ECD were expressed in 293T cells and then purified. Biotinylated ACE2-ECD (ACE2-ECD-Biotin) binds to streptavidin-coated Alpha donor beads and His-tagged S-RBD (S-RBD-His) binds to anti-His-conjugated AlphaLISA acceptor beads. When S-RBD binds to ACE2-ECD, the two beads come into close proximity. Upon illumination at 680 nm, the donor beads generate singlet oxygen molecules that diffuse to acceptor beads and transfer energy to thioxene derivatives in the acceptor beads resulting in light emission at 520-620 nm. The results showed that the incubation of ACE2-ECD-Biotin with S-RBD-His produced very strong AlphaScreen signal, and the signal decreased to the basal level in the absence of either of the two proteins (Fig. 1b). To confirm the specificity of this AlphaScreen system, we replaced S-RBD-His with His-tagged extracellular domains of other membrane proteins, including mucosal vascular addressin cell adhesion molecule 1 (MAdCAM-1) and vascular cell adhesion molecule 1 (VCAM-1). Co-incubation of MAdCAM-1-His or VCAM-1-His with ACE2-ECD-Biotin did not generate AlphaScreen signal, indicating that the system detects S-RBD—ACE2 interaction specifically (Fig. 1c).

**Fig.1.**
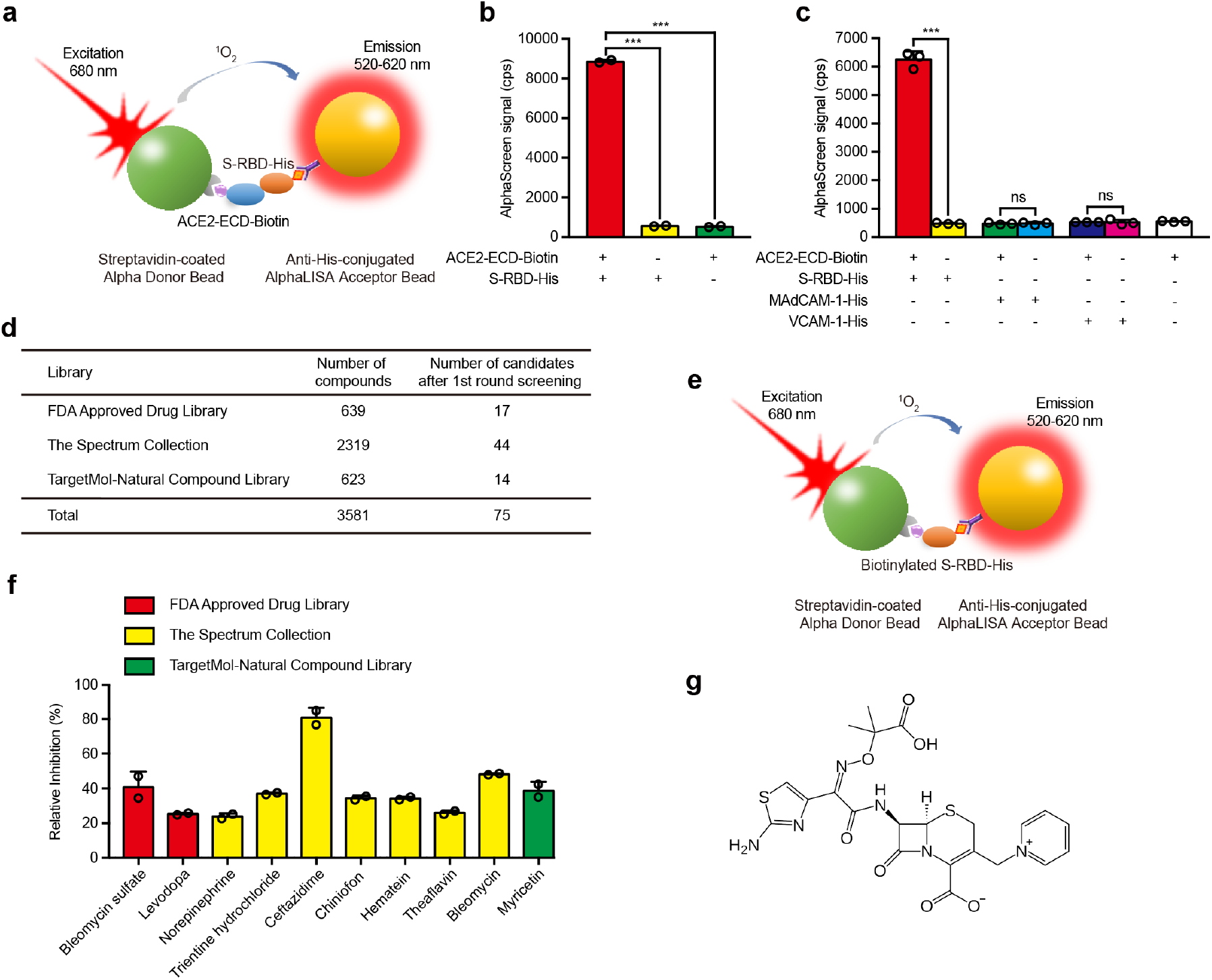
Screening of small molecule compounds that specifically block the interaction between S-RBD and ACE2. **a**, Schematic diagram of AlphaScreen system to detect the interaction between S-RBD and ACE2-ECD. The donor and acceptor beads are coated with streptavidin and anti-His monoclonal antibody, respectively. **b**, The interaction between S-RBD and ACE2-ECD was monitored using AlphaScreen system. **c**, Comparison of the AlphaScreen signal of S-RBD-His, MAdCAM-1-His and VCAM-1-His proteins in the presence or not of ACE2-ECD-Biotin in AlphaScreen system. **d**, Libraries used in AlphaScreen-based high-throughput system and 75 candidates were identified from 3581 compounds in positive selection. The inhibition rate was calculated by the decrease of AlphaScreen signal of each compound compared with that of DMSO vehicle control group. **e**, Schematic diagram of negative selection using AlphaScreen system. Biotinylated S-RBD-His simultaneously links streptavidin-coated donor bead and anti-His-conjugated acceptor bead together to generate AlphaScreen signal directly. **f**, Relative inhibition of 10 candidate compounds on S-RBD–ACE2 interaction using AlphaScreen system. The relative inhibition rate was calculated by subtracting the inhibition rate in negative selection from that in positive selection. **g**, Molecular structure of ceftazidime. Data represent the mean ± SEM (n ≥ 2) in **b**, **c** and **f**. *** p < 0.001, ns: not significant (Student’s t test).

### High-throughput screening of small molecules blocking S-RBD–ACE2 interaction

Next, we used the AlphaScreen-based high-throughput system to screen small molecules that block S-RBD–ACE2 interaction. A total of 3581 small molecule compounds with known molecular structures from FDA Approved Drug Library, Spectrum Collection and Targetmol-Natural Compound Library were assessed (Fig. 1d). The assay was conducted at a final compound concentration of 10 μM and the interaction between S-RBD-His (0.1 μM) and ACE2-ECD-Biotin (0.2 μM) was analyzed. After first round screening, 75 candidate compounds were identified, which showed inhibitory effect on S-RBD–ACE2 interaction (Fig. 1d). All these compounds showed over 45% inhibition rate according to the changes in AlphaScreen signal. To exclude the interference of the compounds to the AlphaScreen system per se, we designed a negative selection system in which the biotinylated S-RBD-His links streptavidin-coated Alpha donor bead and anti-His-conjugated AlphaLISA acceptor bead together to generate AlphaScreen signal directly (Fig. 1e). After the negative selection, 10 compounds, including bleomycin sulfate, levodopa, norepinephrine, trientine hydrochloride, ceftazidime, chiniofon, hematein, theaflavin, bleomycin and myricetin, from the 75 candidate compounds were validated to inhibit S-RBD—ACE2 interaction effectively. Among the 10 compounds, ceftazidime was the most potent inhibitor which showed a relative inhibition rate of 80.7% (Fig. 1f). Thus, ceftazidime was selected for further investigation considering the best inhibitory effect on S-RBD—ACE2 interaction, the anti-inflammatory effect and the minimal side effect of this drug compared with the other 9 compounds ^14–17^ (Fig. 1g).

### Ceftazidime specifically binds to S-RBD protein

To investigate whether S-RBD or ACE2 is the binding target protein of ceftazidime, we applied a bio-layer interferometry experiment to examine the binding affinity between ceftazidime and S-RBD or ACE2-ECD. Along with the elevated concentration of ceftazidime, this compound showed increased binding to S-RBD protein with an K_D_ value of 260 ± 38 μM (Fig. 2a). Notably, ceftazidime hardly dissociated from S-RBD, indicating a strong and stable interaction between ceftazidime and S-RBD. By contrast, ceftazidime and ACE2-ECD showed no specific binding signal (Fig. 2b). Thus, ceftazidime binds to S-RBD specifically.

**Fig.2.**
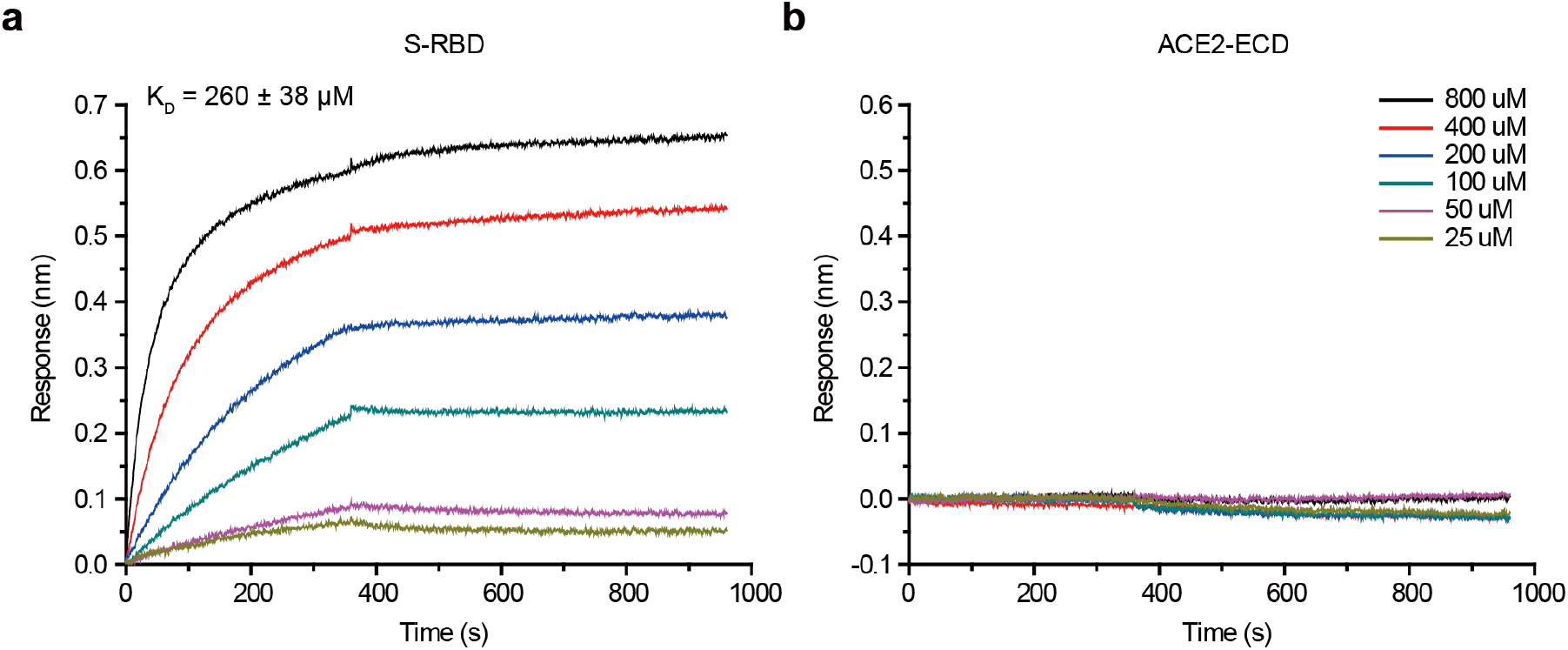
Ceftazidime specifically binds to SARS-CoV-2 S-RBD. Binding profiles of ceftazidime to S-RBD and ACE2-ECD were measured by bio-layer interferometry in an Octet RED96 instrument. The biotin-conjugated S-RBD or ACE2-ECD was captured by streptavidin that was immobilized on a biosensor and tested for binding with gradient concentrations of ceftazidime.

### Ceftazidime inhibits S-RBD binding to human pulmonary alveolar epithelial cells

Lung is the main organ infected by SARS-CoV-2, which cause severe acute respiratory syndrome (SARS) ^18,19^. Therefore, we examined the inhibitory effect of ceftazidime on the binding of S-RBD protein to human pulmonary alveolar epithelial cells (HPAEpiC) which express ACE2. Addition of 100 μM ceftazidime into the soluble S-RBD binding assay system led to a significantly decrease in the S-RBD binding signal (Fig. 3a), demonstrating the efficient inhibition on S-RBD binding to HPAEpiC cells by ceftazidime. Further analysis showed an IC_50_ of 39.90 ± 1.11 μM (Fig. 3b).

**Fig.3.**
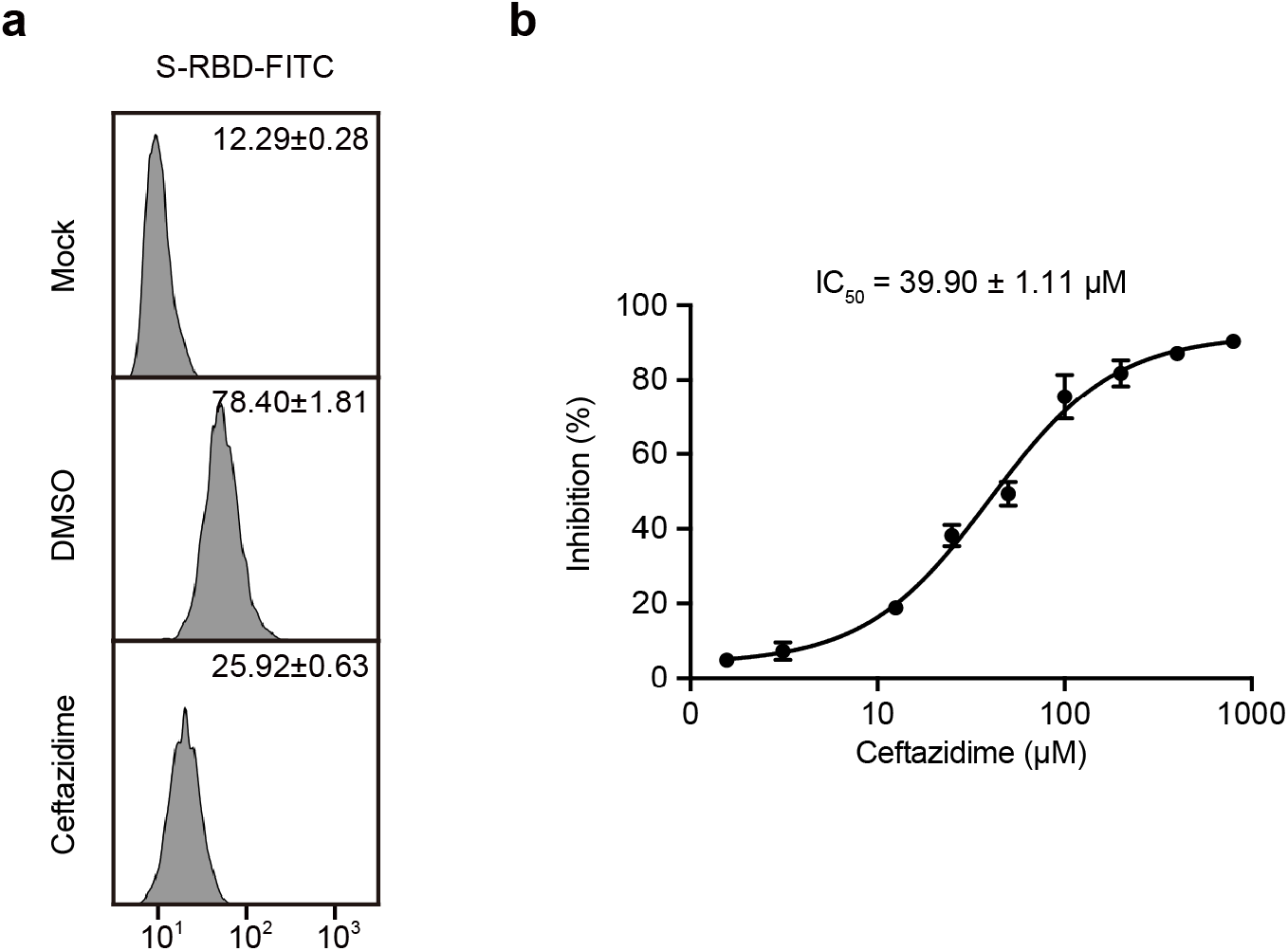
Ceftazidime inhibits S-RBD binding to HPAEpiC cells. **a**, Soluble S-RBD binding to HPAEpiC cells was examined by flow cytometry analysis. Mock, cells were incubated with FITC-conjugated goat anti-human IgG; DMSO, vehicle control; Ceftazidime, 100 μM ceftazidime in DMSO. Numbers within the panel showed the specific mean fluorescence intensities. **b**, The inhibitory effect of ceftazidime on the binding of S-RBD protein to HPAEpiC cells. Cells were treated with different concentrations of ceftazidime. The inhibition rate was calculated by the decrease of the mean fluorescence intensity of each group compared with that of DMSO vehicle control group. IC_50_ was indicated in the graph. One representative result of three independent experiments is shown in **a**. Data represent the mean ± SEM (n = 2).

### Ceftazidime inhibits SARS-CoV-2 pseudovirus infection

Pseudovirus has the similar infectivity compared with authentic virus, thus has been widely applied to carry out the research on the intrusion mechanism of virus with high infectivity and pathogenicity ^4^. To evaluate the inhibitory effect of ceftazidime on the entry of SARS-CoV-2 pseudovirus into 293T cells overexpressing human ACE2, we added ceftazidime into the SARS-CoV-2 pseudovirus infection assay system at a series of concentrations. The results showed that ceftazidime efficiently inhibited SARS-CoV-2 pseudovirus cell entry *in vitro* and the IC_50_ was 113.24 ± 1.23 μM (Fig. 4). Of note, ceftazidime showed no detectable cytotoxicity at a high concentration of 400 μM, indicating its safety for clinical usage (Fig. 4). Thus, ceftazidime has both anti-bacterial and anti-SARS-CoV-2 effects and should be considered as the first-line antibiotics used for the treatment of COVID-19.

**Fig.4.**
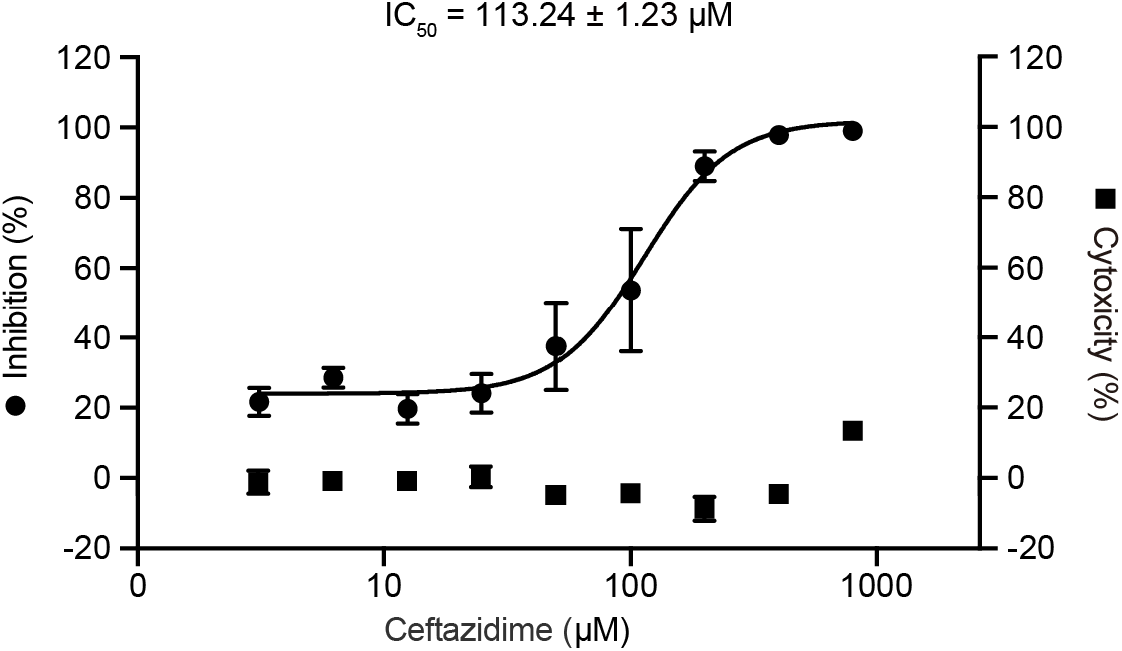
Ceftazidime inhibits SARS-CoV-2 pseudovirus infection. Inhibition of luciferase-encoding SARS-CoV-2 typed pseudovirus entry into ACE2-expressing 293T cells by ceftazidime. Cells were treated with different concentrations of ceftazidime. IC_50_ was indicated in the graph. The cytotoxicity of ceftazidime to 293T cells was determined by the CCK8 assay. Data represent the mean ± SEM (n ≥ 2).

## DISCUSSION

Up to date, studies on COVID-19 therapeutics development mostly focus on screening and validation of neutralizing antibodies and the development of vaccine, both require a relatively long time for the effectivity validation and safety assessment. An effective drug with minimal side effect for COVID-19 treatment is eagerly needed. Therefore, repurposing of FDA-approved drugs with minimal known side effects will accelerate the deployment of effective and affordable therapies for COVID-19. In this study, we found that ceftazidime, an antibiotic used for the treatment of pneumonia, has strong inhibitory effect on SARS-CoV-2 pseudovirus cell entry *in vitro* by inhibiting the S-RBD–ACE2 interaction. It is noteworthy that ceftazidime has anti-SARS-CoV-2 cell entry and anti-bacterial infection dual functions with little known side effects, which make is ready for immediate preclinical and clinical trials for the COVID-19.

Ceftazidime has been widely used in the treatment of bacterial infections. Compared with other compounds that we identified showing inhibition on S-RBD–ACE2 interaction, ceftazidime has less toxicity and side effects. Except for the patients with allergic history of cephalosporins, most COVID-19 patients can be treated with this drug. Ceftazidime has been clinically used as a drug for the treatment of bacterial pneumonia and the blood concentration of ceftazidime can reach over 300 μM. At this concentration, ceftazidime showed an 96% inhibition of SARS-CoV-2 pseudovirus infection *in vitro*, indicating its strong potency to inhibit cell entry of SARS-CoV-2.

Cephalosporins have many derivatives which share the similar core structure but have different side-chain modifications. We have also compared the inhibitory effects of 14 different cephalosporins, including ceftazidime, cephradine, cefazolin, cephalexin, cefuroxime, cefamandole, cefuroxime axetil, cefotaxime, ceftriaxone, cefoperazone, cefoselis, cefepime, ceftobiprole and ceftaroline. Among all cephalosporins, only ceftazidime showed strong inhibition on the S-RBD–ACE2 interaction. Ceftobiprole and ceftriaxone showed limited inhibitory effects, whereas other cephalosporins have little or no inhibitory effect on S-RBD–ACE2 binding (Extended Data Fig. 1). These results in combination with a preliminary Structure Activity Relationship (SAR) analysis suggested that the unique moieties in ceftazidime, including 2-aminothiazole, oxime protected with a terminal-exposed isobutyric acid and the positive charged pyridine, might be involved in mediating the binding to S-RBD and eventually blocked the protein interaction between S-RBD and ACE2. Furthermore, our data showed that ceftazidime hardly dissociated from the S-RBD proteins (Fig. 2a), which could be due to the covalent binding of ceftazidime to S-RBD.

**Extended Data Fig. 1.**
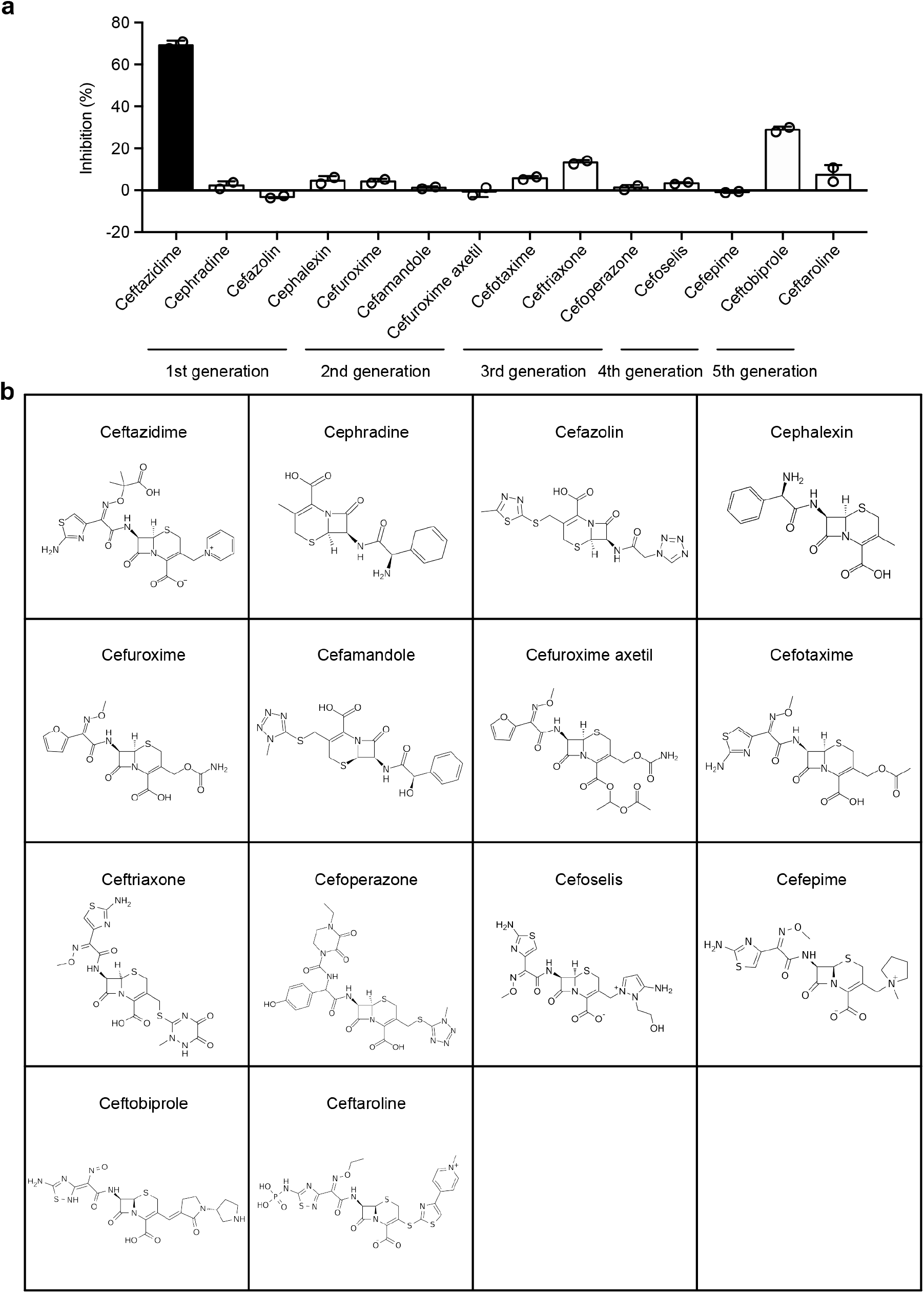
Effect of ceftazidime and the derivatives of cephalosporins on S-RBD–ACE2 interaction. **a**, Inhibition of S-RBD–ACE2 interaction by ceftazidime and the derivatives of cephalosporins was analyzed using AlphaScreen system. **b**, Molecular structures of ceftazidime and the derivatives of cephalosporins. Data represent the mean ± SEM (n = 2) in **a**.

In summary, we identify ceftazidime as a potent inhibitor of SARS-CoV-2 pseudovirus cell entry *in vitro* by binding to S-RBD and consequently blocking S-RBD interaction with ACE2. Since ceftazidime is a drug clinically used for the treatment of pneumonia with affordable price and minimal side effects compared with other antiviral drugs, ceftazidime should be considered as the first-line antibiotics used for the treatment of COVID-19, which deserves the immediate preclinical and clinical trials. Optimization of the molecular structure of ceftazidime may further improve the inhibitory effect of this drug on SARS-CoV-2 infection.

## ACKNOWLEDGEMENTS

This work was supported by grants from the National Natural Science Foundation of China (31525016, 31830112), Personalized Medicines-Molecular Signature-based Drug Discovery and Development, the Strategic Priority Research Program of the Chinese Academy of Sciences (XDA12010101), the CAS/SAFEA International Partnership Program for Creative Research Teams, the Youth Innovation Promotion Association of the Chinese Academy of Sciences and the Young Elite Scientist Sponsorship Program by CAST (2019QNRC001). The authors gratefully acknowledge the support of SA-SIBS scholarship program. We thank Prof. Haitao Yang and Dr. Zhenming Jin from ShanghaiTech University for providing the plasmids expressing spike protein and ACE2, and Dr. Xu Han from Shanghai Institute of Materia Medica, Chinese Academy of Sciences for helpful discussion on the structure of the compounds.

## AUTHOR CONTRIBUTIONS

C.D.L. and J.F.C. designed experiments. C.D.L., Y.L., M.Y.Y., M.W.H., C.L., H.D., X.C.P. and Y.T.W. performed experiments and analyzed data. X.Y.X. and C.Q.X. provided SARS-CoV-2 pseudoviruses. C.D.L., Y.L. and J.F.C. interpreted results. The manuscript was drafted by C.D.L., Y.L. and edited by J.F.C.

## METHODS

### Protein expression and purification

Recombinant SARS-CoV-2 S-RBD fused with Fc/His tag (S-RBD-His) was produced in 293T cells and purified with Protein A Agarose (Thermo Fisher Scientific). Recombinant human ACE2-ECD fused with Flag/His tag was produced in 293T cells and purified with anti-DYKDDDDK G1 Affinity Resin according to the manufacturer’s instructions (GenScript).

### AlphaScreen

AlphaScreen assays were performed in Costar 384-well microplates in a 20 μl final reaction volume. Streptavidin-coated Alpha donor beads or anti-His-conjugated AlphaLISA acceptor beads (PerkinElmer) were used at a final concentration of 10 μg/mL per well. The assays were performed in PBS buffer (155 mM NaCl, 1.06 mM KH_2_PO_4_, 2.97 mM Na_2_HPO_4_, pH 7.4) and 0.1% BSA. 5 μl S-RBD-His (final concentration 0.1 μM) and 5 μl ACE2-ECD-Biotin (final concentration 0.2 μM) were pre-incubated with compounds at a final concentration of 10 μM for 0.5 h at 4 °C. Then donor beads and acceptor beads were added into the reaction in dark for 2 h, at room temperature. Laser excitation was carried out at 680 nm, and readings were performed at 520 to 620 nm using the EnVision (PerkinElmer) plate reader.

### Flow cytometry

0.1 μM S-RBD-Fc/His was pre-incubated with 5 μg/mL FITC-conjugated goat anti-human IgG in 50 μL of PBS and then incubated with HPAEpiC cells for 30 min at room temperature. Cells were washed twice before flow cytometry analysis. Cells were incubated with FITC-conjugated goat anti-human IgG merely as a control.

### Pseudovirus infection assay

SARS-CoV-2 pseudoviruses were produced as previously described ^20^. The pseudoviruses were diluted in complete DMEM mixed with an equal volume (50 μl) of diluted DMSO or ceftazidime, and then incubated at 37 °C for 1 h. The mixture was transferred to 293T expressing human ACE2 stable cell line cells. The cells were incubated at 37 °C for 48 h, followed by lysed with Bio-Lite Luciferase Assay Buffer and tested for luciferase activity (Vazyme). The percent neutralization was calculated by comparing the luciferase value of ceftazidime treatment group to that of DMSO control.

### Bio-layer Interferometry (BLI) Experiment

The BLI experiment was performed using an Octet Red 96 instrument (ForteBio, Inc.). Briefly, biotinylated S-RBD or ACE2-ECD (50 μg/ml) were immobilized on streptavidin (SA) biosensors and then incubated with gradient concentrations of ceftazidime in kinetics buffer (PBS and 0.02% Tween-20). The association and dissociation steps were set to 360 s and 600 s. The KD value of the S-RBD binding affinity for ceftazidime was calculated from all the binding curves based on their global fit to a 1:1 Langmuir binding model with an R^2^ value of ≥ 0.95. Binding experiments were performed at 25 °C. Data were analyzed using Octet Data Analysis Software 9.0 (ForteBio, Menlo park, CA, USA).

## QUANTIFICATION AND STATISTICAL ANALYSIS

Statistical significance was determined by Student’s t test (GraphPad, version 5.01). The resulting p values are indicated as follows: ns, not significant; *, p < 0.05; **, p < 0.01; ***, p < 0.001. Data represent the mean ± SEM of at least two independent experiments.

